# Tissue-Targeted Prime/Pull/Keep Therapeutic Herpes Simplex Virus Vaccine Protect Against Recurrent Ocular Herpes Diseases in HLA-A*0201 Transgenic Rabbits

**DOI:** 10.1101/2024.09.29.615649

**Authors:** Aziz A Chentoufi, Swayam Prakash, Hawa Vahed, Afshana Quadiri, Lbachir BenMohamed

## Abstract

Herpes Simplex Virus type 1 (HSV-1) continues to be one of the most prevalent viral infections globally, with approximately 3.72 billion individuals affected worldwide. A clinical herpes vaccine is still lacking. In the present study, a novel prime/pull/keep vaccine was tested in Human Leukocyte Antigen transgenic rabbit model of ocular herpes (HLA-A*0201 Tg rabbit). Ten asymptomatic (ASYMP) CD8+ T cell peptide epitopes and 3 CD4+ T-cell epitopes were selected from the HSV-1 glycoproteins D and B (gD and gB), viral tegument proteins (VP11/12 and VP13/14) and the DNA replication binding helicase (UL9), all preferentially recognized by CD8^+^ and CD4^+^ T-cells from “naturally protected” HSV-1-seropositive healthy ASYMP individuals (who never had recurrent corneal herpetic disease). HLA Tg rabbits were ocularly infected with HSV-1 then during latency at day 30 post-infection, the rabbits were ocularly vaccinated with a recombinant neurotropic AAV8 vector (10^7^GC/ eye) encoding for the 10 CD8+ T cell peptide and 4 CD4+ T cells peptide (prime), T cell-attracting CXCL-11 (Pull) and T-cell keeping IL-2/IL-15 cytokines (keep). The rabbits were followed up for eye disease and viral loads in tears for 28 days. The frequency, function and protective efficacy of HSV-specific CD8^+^ T cells induced by the prime/pull/keep vaccine were assessed in the trigeminal ganglia (TG), cornea, Spleen, and peripheral blood. Compared to the mock group (unvaccinated), the peptides/CXCL11/IL-2/IL-15 vaccine generated frequent resident CD8^+^ T cells that infiltrated the TG. CD8^+^ T cells mobilization and retention into TG of prime/pull/keep vaccinated rabbits was associated with a significant reduction in corneal herpes infection and disease after an ocular HSV-1 challenge (McKrae). These findings draw attention to the novel prime/pull/keep therapeutic vaccine strategy to mobilize and retain anti-viral T cells into tissues protecting them against herpetic infection and disease.

**IMPORTANCE:** There is an urgent need for a vaccine against widespread human herpes simplex virus infections. The present study demonstrates that immunization of humanized HLA-A*0201 transgenic rabbits with CD8+ and CD4+ T-cell epitope peptides (prime)/ CXCL11 (pull)/ IL-2/IL-15 (Keep) AAV8-based vaccine triggered mobilization ad retention of HSV-1-specific CD8^+^ T cells locally in the cornea and TG, the sites of acute and latent herpes infections. Mobilization and retention of antiviral CD8^+^ T cells into cornea and TG of HSV-1 infected rabbits that received the prime/pull/keep vaccine was associated with protection against ocular herpes infection and disease. These results highlight the importance of the prime/pull/keep vaccine strategy to bolster the number and function of protective CD8^+^ T cells within infected tissues.

## INTRODUCTION

Herpes Simplex Virus type 1 (HSV-1) continues to be one of the most prevalent viral infections globally, with approximately 3.72 billion individuals affected worldwide (1, 2). HSV-1 associated ocular herpetic infection range from asymptomatic to symptomatic clinical conditions including blepharitis, conjunctivitis, neovascularization, disciform stromal edema, and the vision-threatening condition herpetic stromal keratitis (HSK) (3–10). More than 450,000 individuals in the United States alone, have suffered from blinding ocular herpetic disease, necessitating medical intervention such as antiviral drug treatment, and corneal transplantation (11–13). Despite current antiviral therapies, such as Acyclovir and its derivatives that merely reduce the symptoms of herpetic disease by about 45% Kuo, 2014 #75932}, HSV-1 ocular infection continue to rise globally and the development of an effective vaccine would represent a critical advancement, offering a more sustainable and cost-effective method to combat corneal herpetic infections and disease (reviewed in (16)).

Evidence from human cadaveric brain studies (14–16) has shown that CD8+ T cells residing in both the trigeminal ganglia (TG) and cornea play an essential role in controlling reactivation of latent HSV-1 from sensory neurons, thereby preventing corneal herpetic infections and disease (17, 18). However, herpes vaccine clinical trials, which employed recombinant glycoproteins B and D (gB and gD), failed to protect humans from herpetic disease despite generating high levels of HSV-specific neutralizing antibodies (19, 20).These disappointing results highlight the importance of inducing strong T-cell responses, in addition to antibody-mediated immunity, to achieve protective immunity against ocular herpes. For an effective T cell mediated immunity against viral pathogens, it is fundamentally important for these T cells to migrate to the site of infection and an optimal number of memory resident T-cells to be established.

The migration of HSV-specific CD8+ T cells to key anatomical sites of latent infection, such as the TG, is thought to be tightly regulated by interactions between chemokines and their corresponding chemokine receptors, a process that can be influenced by the vaccine strategy employed (21–25). Increasing evidence supports the notion that CD8+ T cells specific to HSV epitopes possess intrinsic characteristics that guide their migration to the cornea and TG, which are the primary sites of acute and latent HSV-1 infections respectively. However, the TG is recognized as an immunologically restrictive environment, meaning it is not easily accessible to CD8+ T cells that are activated by a conventional parenteral vaccine and subsequently enter the circulation (26). This is thought to be due to low levels of T cell-attracting chemokines, such as CXCL9, CXCL10, CXCL11, and CCL5, in non-infected TG tissue, which limits the migration of sufficient antiviral CD8+ T cells from the bloodstream into the TG.

In this study, we proposed that a prime/pull/keep AAV8-based vaccine strategy, designed to both “prime” functional antiviral CD8+ T cells in peripheral tissues and “pull” and “keep” them in the infected cornea and TG, would lead to a significant reduction in corneal HSV-1 infection and disease. To evaluate this hypothesis, we utilized the recently developed “humanized” HLA-A*0201 transgenic rabbits (HLA Tg rabbits), which represent the most reliable small animal model for studying ocular herpes infection and associated diseases (17, 18, 27–29). Our results showed that immunizing HLA-A*0201 Tg rabbits with a topical ocular application of a recombinant neurotropic AAV8 vector encoding for a combination of immunodominant HSV-1 CD8+ T-cells and CD4+ T-cells epitopes (prime) selected based on their preferential recognition by CD8+ and CD4+ T cells from HSV-1 seropositive asymptomatic individuals who are naturally protected from disease and the chemokine CXCL-11 (pull) and IL-2/IL-15 cytokines (Keep), resulted in: (i) robust induction of HSV-specific CD8+ T cells in the TG, characterized by their phenotype of resident memory T-cell and (ii) enhanced local recruitment of CD8+ T cells to the TG. Importantly, this mobilization of antiviral CD8+ T cells correlated with a significant reduction in ocular HSV-1 infection and decreased severity of herpetic recurrent corneal disease.

These preclinical findings in the HLA-A*0201 Tg rabbit model, a well-established model of ocular herpes infection, strongly suggest that the prime/pull/keep vaccine strategy—based on HSV-1 human epitope peptides combined with CXCL11 and IL-2/IL-15 cytokines—has the potential to provide robust, safe, and protective immunity. This approach should be considered in the future development of a clinical vaccine for ocular herpes, as it offers a promising solution for controlling both HSV-1 infection and herpetic eye disease.

## MATERIALS AND METHODS

### HLA-A*02:01 transgenic rabbits

HLA Tg rabbits were derived from New Zealand White rabbits (18). The HLA Tg rabbits retain their endogenous rabbit MHC locus and express human HLA-A*0201 under the control of its normal promoter (18). Prior to this study, the expression of HLA-A*0201 molecules on the PBMC of each HLA Tg rabbit was confirmed by FACS analysis. In brief, PBMCs were stained with 2 μl anti–HLA-A2 mAb, BB7.2 (BD Pharmingen, San Diego, CA), at 4°C for 30 min, washed and analyzed by flow cytometry using a LSRII (Becton Dickinson, Mountain View, CA). The acquired data were analyzed with FlowJo software (TreeStar, Ashland, OR). All rabbits used in these studies had a similarly high level of HLA-A*02:01 expression (>90%). This eliminated any potential bias due to the variability of HLA-A*0201 molecule levels in different animals. New Zealand White rabbits (non-Tg control rabbits), purchased from Western Oregon Rabbit Co., were used as controls. All rabbits were housed and treated in accordance with ARVO (Association for Research in Vision and Ophthalmology), AAALAC; Association for Assessment and Accreditation of Laboratory Animal Care, and NIH (National Institutes of Health) guidelines.

### Peptide synthesis

CD4+ and CD8+ T cell epitope peptides from the HSV-1 glycoproteins D and B (gD and gB), viral tegument proteins (VP11/12 and VP13/14) and the DNA replication binding helicase (UL9) proteins were synthesized by 21^st^ Century Biochemicals (Marlboro, MA). All peptides (CD8+ T cell epitopes: *gD_53-61_, gD_278-296_, gB_183-191_, gB_342-350_, gB_561-569_,VP11/12_220-228_, VP11/12_702-710_, VP13/14_504-512_, VP13/14_544-552_, UL9_196-204_ and CD4+ T-cell epitopes: gB_166-180_, VP11/12_129-143_, gD_49-82_)* were HPLC purified to a purity of 95% to 98% as determined by reversed-phase high-performance liquid chromatography (Vydac C18) and mass spectroscopy (Voyager MALDI-TOF System). Stock solutions were made at 1 mg/ml in PBS. All peptides were aliquoted and stored at −20°C until assayed.

### Design and construction of AAV8 vector expressing CD8+ and CD4+ T-cell epitopes, CXCL11 chemokine and IL-2/IL-15 under neurotropic CamKIIα promoters

Two prototype PPK vaccine molecules. The single AAV8-vectors expressed 5 pairs of CD4+-CD8+ T cell epitopes together with CXCL11 and IL-2 or IL-15. Both candidate vaccines expressed the same 5 CD8+ epitopes in pairs with the CD4+ T helper epitope. The PPK vaccines were made by vector Biolabs, Inc., where rabbit CXCL11 is co-expressed by an AAV8 vector in tandem with the T cell epitopes, under the control of the CamKIIα neuron-specific promoter.

### Prime/pull/keep vaccination after ocular herpes challenge

HLA-A*0201 Tg rabbits (8-10 weeks) with similar high expression of HLA-A*02:01 molecules (>90%) were used, as described below. Groups of age-matched HLA-A*02:01 rabbits (n=2 groups; each group has 5 rabbits) were ocularly infected with HSV-1 (5×10^5^ pfu). On day 30, half of the rabbits were left untreated (mock vaccinated), and the other half received a topical ocular treatment with 10^10^ viral genome copies (GC) of a rAAV8-CamKIIα-NP7-CamKIIα−CXCL11-CamKIIα−IL-2/−CamKIIα−IL15 vectors.

### Herpes simplex virus production

HSV-1 (strain McKrae) was used in this study. The virus was triple plaque purified and prepared as previously described (23, 24) using rabbit skin cell monolayers grown in MEM (1×) containing 10% FBS (HI), 2mML-glutamine, 2.5ug/ml amphotericin, and 5% penicillin-streptomycin solution (from a stock of 10,000 IU penicillin and 10,000 μg/ml streptomycin).

### Ocular infection of rabbits with HSV-1

Without making any corneal sacrifice, rabbits were ocularly (both eyes) infected by dropping 5 μl HSV-1 (2 × 10^5^ PFU) strain McKrae suspended in culture medium on day 0(8).

### Rabbit corneal disease clinical scores

Rabbits were examined for ocular disease and survival for 30 days after the challenge. The ocular disease was determined by a masked investigator using fluorescein staining and slit lamp examination before challenge and on days, 7, 14, 21 and 28 thereafter. A standard 0-4 scale: 0, no disease; 1, 25%; 2, 50%; 3, 75%; and 4, 100% staining, was used.

### Quantification of ocular infectious virus

Tears were collected from both eyes using a Dacron swab (type 1; Spectrum Laboratories, Los Angeles, CA) on days 7, 14, 21, and 28 post vaccination. Individual swabs were transferred to a 2-ml sterile cryogenic vial containing 500 μl culture medium and stored at −80°C until use. The HSV-1 titers in tear samples were determined by standard real-time PCR based on previously described reaction conditions (30).

### Preparation of Rabbit cell suspensions from TG and spleen

Rabbits were euthanized TG, and spleen were isolated and finely minced using dissection scissors. Then digested in DMEM– 5% fetal bovine serum (FBS) containing collagenase I (Life Technologies, Carlsbad, CA), as we previously described (5, 16, 17, 37). The digested tissue suspension was passed All samples were placed at 37°C in a shaker at 250rpm and incubated for 1hr (TG) and 15min (spleen) and passed through a 70-μm nylon cell strainer followed by a 40-μm nylon cell strainer. The cell suspensions were centrifuged and resuspended in complete media then lymphocyte population were isolated on Percoll gradient 40%. Cell suspensions were spun down at 1,400 rpm for 5 min at 4°C and then washed and suspended in fluorescence-activated cell sorter (FACS) buffer (phosphate-buffered saline [PBS]–0.01% NaN3–0.1% bovine serum albumin [BSA], 2 mM EDTA) for FACS acquisition and analysis.

### Rabbit peripheral blood mononuclear cells isolation

10 mL of blood was drawn from each rabbit into a yellow-top Vacutainer^®^ Tubes (Becton Dickinson, USA). Sera were isolated by centrifugation for 10 min at 800g. PBMCs were isolated by gradient centrifugation using leukocyte separation medium (Cellgro, USA). The cells were washed in PBS and re-suspended in complete culture medium consisting of RPMI-1640 medium containing 10% FBS (Bio-Products, Woodland, CA, USA).

### Flow cytometry assays on rabbit T cells

Cell suspensions from TG, Blood and spleen were analyzed by flow cytometry after staining with HLA-tetramers-PE and panels of anti-Rabbit CD8, CD4, CD69, CD62L, CD103 and CD44. The following anti-rabbit antibodies were used: anti-CD8 (clone MCA1576F, Serotec) FITC, -CD4 (clone KEN-4) FITC, -CD44 (clone IM7) PE-Cy7, -CD62L (clone DREG-56) Alexa Flour 700, -CD69 (clone FN50) APC/Cy7, -CD103 (clone LF61) PercP.Cy5.5. For surface stain, panels of mAbs against various cell markers were added to a total of 1X10^6^ cells in 1X PBS containing 1% FBS and 0.1% sodium azide (FACS buffer) for 45 min at 4°C. Cells were washed again with Perm/Wash and FACS Buffer and fixed in PBS containing 2% paraformaldehyde (Sigma-Aldrich, St. Louis, MO). For each sample, 500,000 total events were acquired on the BD LSRII. Ab capture beads (BD Biosciences) were used as individual compensation tubes for each fluorophore in the experiment. To define positive and negative populations, we employed fluorescence minus controls for each fluorophore used in this study when initially developing staining protocols. In addition, we further optimized gating by examining known negative cell populations for background level expression.

### Rabbit CD8^+^ T cells tetramer assays

For tetramer-specific CD8^+^ T cell frequencies, TG, Spleen and PBMCs were analyzed for the frequency of CD8^+^ T-cells specific to each of the 10 CD8^+^ T cell epitopes using the corresponding HLA-A2-peptide/Tetramer, provided by the NIH tetramer facility (7, 31). A human beta-2-microglobulin was incorporated in the tetramers, as no rabbit beta-2-microglobulins are currently available. Briefly, the cells were first incubated with 1 μg/ml of each of the three PE-labeled HLA-A2-peptide/Tetramer at 37° C for 30-45 min. The cells were washed twice and stained with panels of anti-Rabbit CD8, CD4, CD62L, CD69, CD103 and CD44. After two additional washings, cells were fixed with 2% formaldehyde in FACS buffer. A total of 500,000 events were acquired by LSRII followed by analysis using FlowJo software. The absolute numbers of individual peptide-specific CD8^+^ T cells were calculated using the following formula: (# of events in CD8^+^/Tetramer^(+)^ cells) x (# of events in gated lymphocytes) / (# of total events acquired).

### Rabbit TG and Cornea Histopathology

Rabbits were euthanized; corneas and TG were harvested, embedded in Tissue-Tek (OCT compound; VWR International, West Chester, PA), and snap-frozen. Approximately 7μm-thick cryo sections were fixed in acetone and stored at −80°C. Sections were H&E stained and mounted. Cell infiltration was examined using a Keyence BZ-X700 microscope at 10X magnification and imaged using z-stack.

### Statistical analyses

Data for each assay were compared by analysis of variance (ANOVA) and Student’s *t*-test using GraphPad Prism version 5 (La Jolla, CA). Differences between the groups were identified by ANOVA and multiple comparison procedures, as we previously described (31). Data are expressed as the mean ± SD. Results were considered statistically significant at *p* < 0.05.

## RESULTS

### 1. A multi-epitope/CXCL11/IL-2/IL-15 (prime/pull/keep) vaccine design

Ten CD8+ T-cell epitope peptides that exhibited high affinities for soluble HLA-A*0201, and 3 CD4+ T-cells epitopes from 5 HSV-1 proteins were selected for the design of PPK vaccine (**Table 1**). We sought to determine whether priming with the mixture of these HSV-1 peptides expressed by AAV-8 encoding for CXCL11 (pull) and IL-2 or IL-15 (keep) would: (i) boost the protective efficacy of HSV-1 epitope specific T cells; and (ii) pull more HSV-1 specific T cells within the cornea and TG, the sites of acute and latent HSV-1 infection in rabbits. Using the optimal CamKIIα promoter we constructed a neurotropic recombinant non-replicating adeno-associated virus type 8 vector (rAAV8) expressing the CD8+ and CD4+ specific T-cell epitope peptides, T-cell attracting CXCL11 chemokine and IL-2/IL-15: rAAV8-multi-epitopes-CamKIIa-CXCL11-CamKIIa-IL-2/IUL-15 (PPK) (**Fig. 1A**).

**Figure 1:**
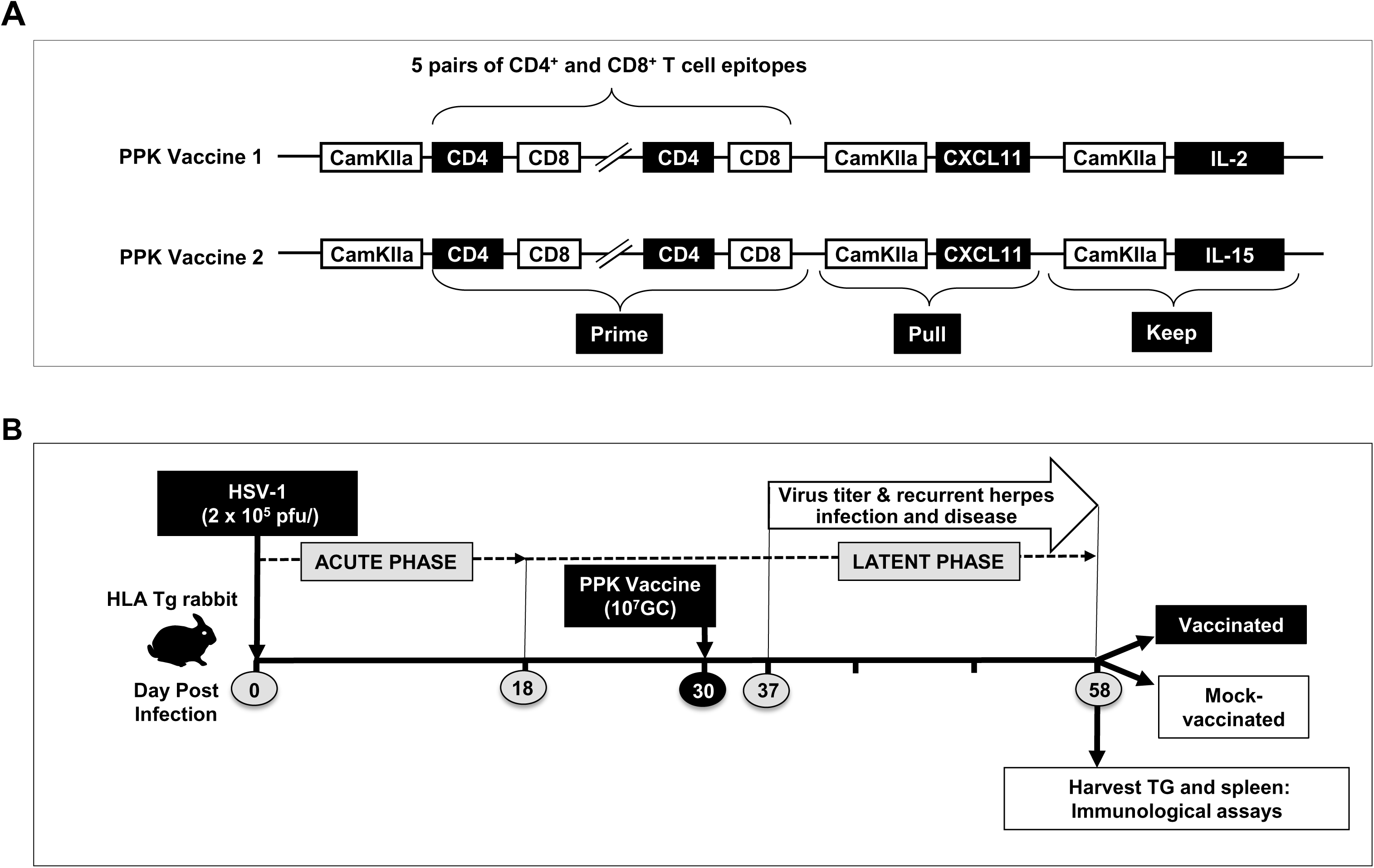
The effect of treatment with a novel prime-pull-keep vaccine approach on herpes disease outcome determined in HLA-A*0201 transgenic rabbits. (**A**) Illustration showing prototype of the two Prime-Pull-Keep (PPK) vaccine approaches. The first approach PPK1 contains multi-epitope herpes vaccine consisting of immunogenic 10 CD8^+^ T cell epitopes, and 3 CD4^+^ T cell epitopes meant to prime the T cells; CXCL11 molecule to pull the T cells; and IL-2 to keep the T cells to protect against herpes simplex virus infection. Whereas the second approach showing vaccine PPK2 contains multi-epitope herpes vaccine consisting of immunogenic 10 CD8^+^ T cell epitopes, and 3 CD4^+^ T cell epitopes meant to prime the T cells; CXCL11 molecule to pull the T cells; and IL-15 to keep the T cells to protect against herpes simplex virus infection. (**B**) Schematic representation of HSV-1 ocular infection of HLA Transgenic rabbits. Rabbits were followed for acute phase of infection for 18 days. Subsequently 30 days post HSV-1 infection the transgenic rabbits were immunized with PPK vaccine at 1 x 10^7^ GC treatment, immunological, virological, and disease analyses in the HLA-A*0201 Tg rabbits ocularly infected. The latent phase of herpes infection was monitoring among the rabbits between Day 37 to Day 58 post HSV-1 infection. Corneal swabs were collected, and disease was monitored during latent phase of infection. On Day 58 the HLA-A*0201 Tg rabbits were euthanized with HSV-1. Trigeminal ganglia and cornea were collected from vaccinated and mock-vaccinated groups of HLA-A*0201 Tg rabbits for immunological and virological assays.

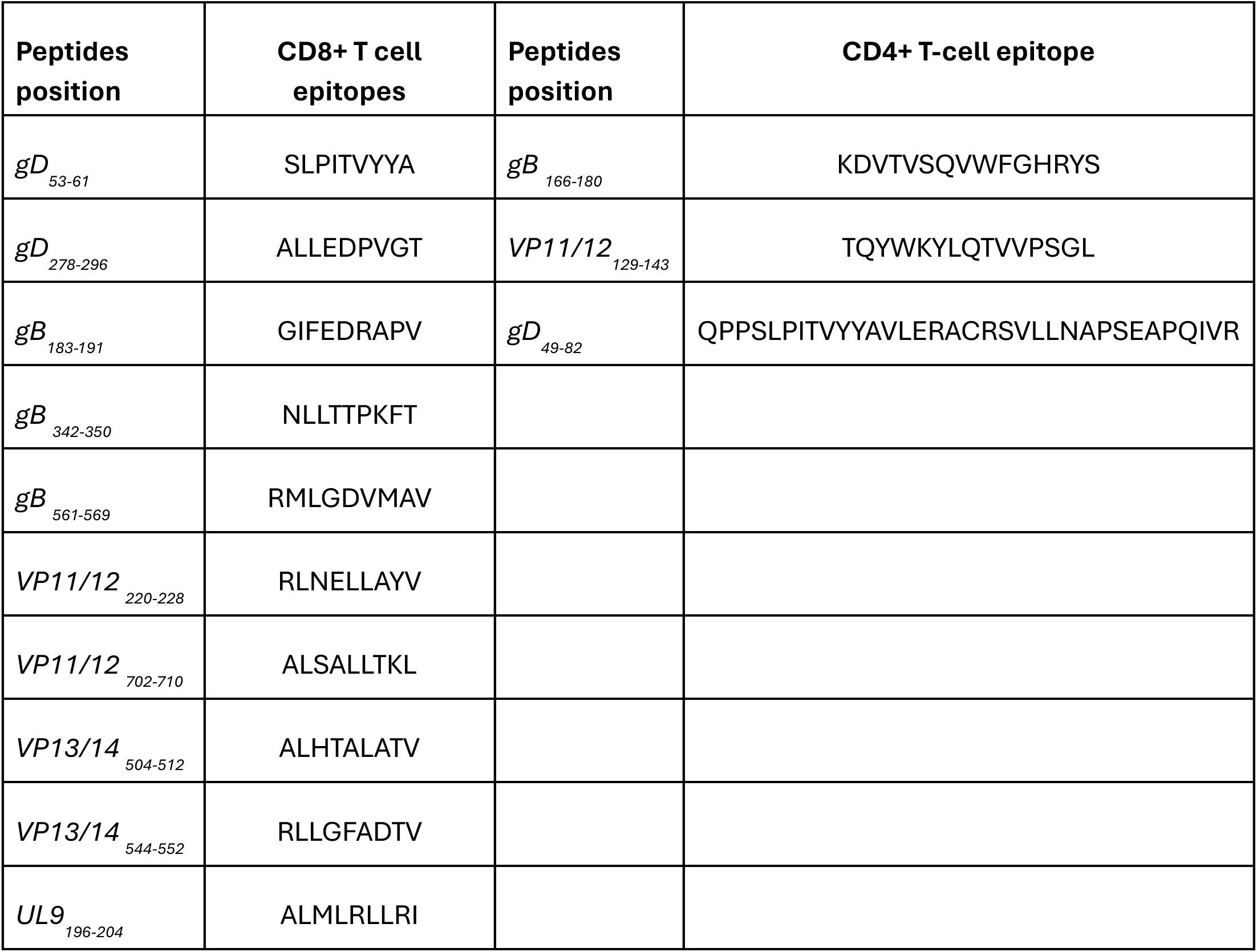

### 2. Selection of Human Leukocyte Antigen (HLA-A*02:01) transgenic rabbits for pre-clinical evaluation of a multi-epitope/CXCL11/IL-2/IL-15 (prime/pull/keep) vaccine against recurrent ocular herpes infection

HLA-A*02:01 positive transgenic rabbit breeders were selected based on their high expression of HLA-A*02:01 molecules, since the expression of the rabbits’ own MHC class I molecules might interfere with the human HLA-A*02:01-restricted responses (material and methods). The high expression of HLA-A*02:01 molecules in the selected HLA Tg rabbits should result in rabbit CD8^+^ T cells using the human HLA-A*02:01 molecules both at the thymus selection and peripheral effector levels (29). All selected HLA Tg rabbits had a similar high expression of HLA-A*02:01 molecules in over 95% of PBMC, as assessed by FACS.

HLA Tg rabbits (n = 12) were ocularly infected with HSV-1 (McKrae, 10^5^ pfu/eye) then vaccinated with the PPK vaccine day 30 postinfection, as shown in **Fig. 1B**. An additional group of HLA Tg rabbits was mock-vaccinated and received an empty AAV8 vector, as control (Mock). Rabbits were followed-up for eye disease and viral loads in the tears for 30 days then sacrificed at day 58 postinfection. The PPK treated group displayed significantly less recurrent corneal herpetic disease (**Fig. 2a**) and virus titers in the eyes (**Fig. 2c**). *In* contrast, the untreated group showed a significantly higher level of virus replication in eyes associated with severe recurrent ocular herpetic disease (**Figs 2a & 2c**). histopathological analysis (**Fig. 2b**) shows that PPK immunized rabbits have a higher number of mononuclear cells in the TGs and the Cornea, suggesting that the immunization with PPK induced higher number of CD4+ and CD8+ T-cells in the site of HSV-1 infection (cornea and TGs).

**Figure 2:**
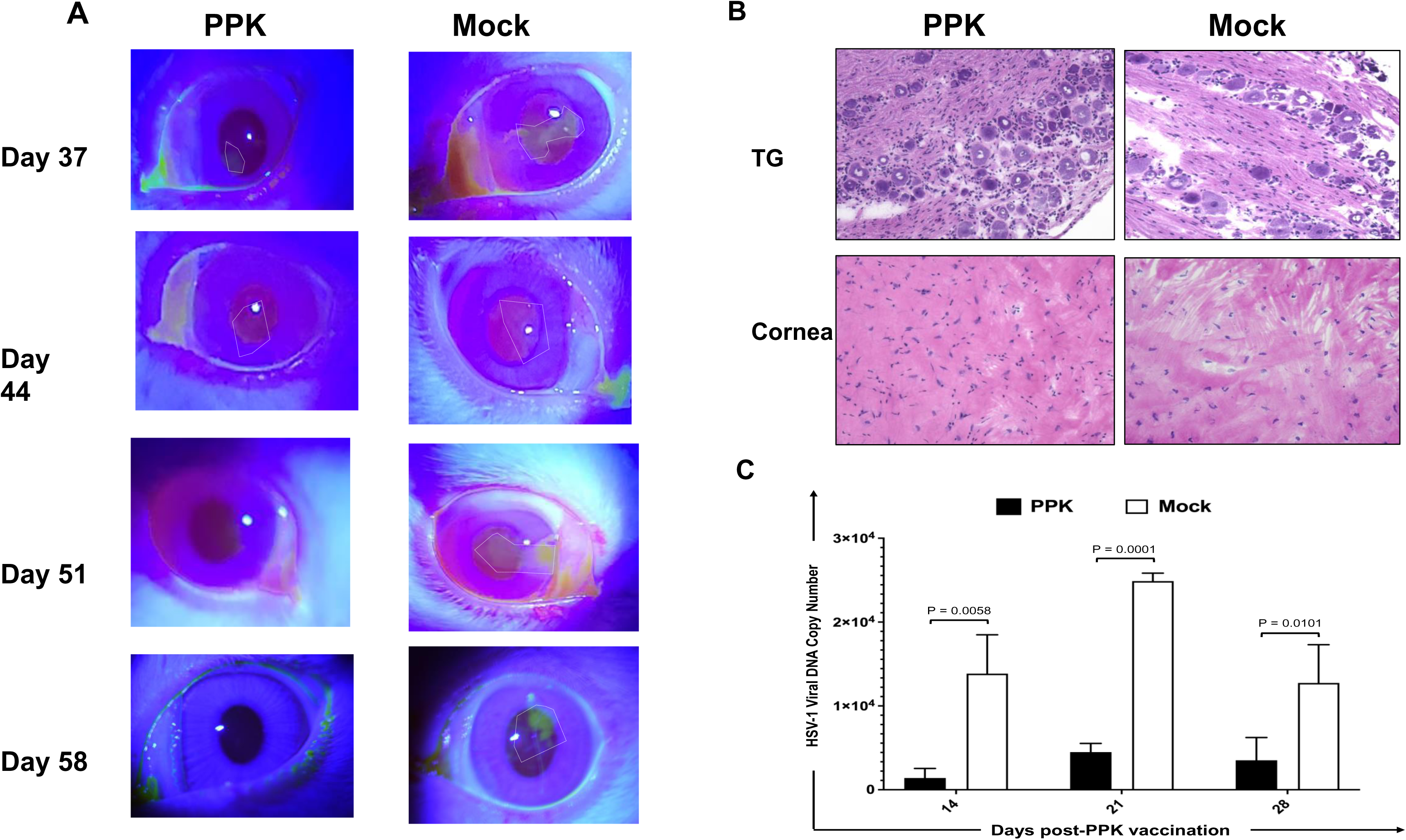
Physical estimation, Histopathological evaluation and HSV-1 Viral DNA copy number showing reduced disease pathology in HLA-A*0201 transgenic rabbits vaccinated with PPK vaccine in comparison to mock-vaccinated rabbits. (**A**) Representative images showing reduced recurrent ocular herpetic disease in HLA-A*0201 Tg rabbits vaccinated with PPK vaccine in comparison to mock-vaccinated HLA-A*0201 Tg rabbits. (**B**) Representative H & E staining images of the trigeminal ganglia and cornea at day 58 *p.i.* of HSV-1 infection in PPK vaccinated and Mock-vaccinated HLA-A*0201 Tg rabbits. Images were taken at 4x magnification. (**C**) Bar diagrams showing comparison of HSV-1 viral titers in the eyes of PPK vaccinated and Mock-vaccinated HLA-A*0201 transgenic rabbits. Infectious virus particles were quantified by quantitative real time PCR (qRT-PCR) from eye swabs of PPK vaccinated and Mock-vaccinated HLA Tg rabbits. The data are representative of one independent experiments and the graphed values and bars represent SD between the two experiments.

### 3. A multi-epitope/CXCL11/IL-2/IL-15 (prime/pull/keep) vaccine bolsters the frequencies of resident memory CD8+ TEM, TCM, and TRM T cells in HLA transgenic rabbits

Since the PPK vaccine enhanced protection as measured by reduced viral loads, and herpetic eye diseases in HLA-transgenic rabbits following HSV-1 infection, we sought to determine whether this protection was associated with increased frequencies of resident memory CD8+ and CD4+T cells in the TGs. Vaccinated/PPK-treated and Mock-vaccinated/untreated rabbits were euthanized on day 58 post-infection, and the frequencies HSV-1 specific CD8+ T-cells of the resident memory CD8+ T cells expressing CD103, CD69; effector memory CD8+ T cells expressing CD62L, and CD44 among total cells [i.e., effector memory (TEM), resident memory (TRM), and central memory (TCM)] were determined by FACS, as described in Materials and Methods. The data in **Fig.3 A** show overall the most dominant epitopes triggered higher number of HSV-1 specific CD8+ T-cells in PPK vaccinated rabbit compared to Mock vaccinated Rabbits. Absolute count of these HSV-1 specific CD8+ T-cells in the TG show that in PPK vaccinated rabbits have 4 to 5 times higher number of HSV-1 Specific CD8+ T-cells in the TG compared to mock rabbits (**Fig. 3B**). The analysis of HSV-specific memory CD8+ T-cells showed a significant increase in memory CD8+ T cells expressing CD44, CD69 and CD62L in PPK vaccinated rabbits (**Fig. 4 A&B**). More interestingly, we found a significant increase in CD103+CD8+ T cells, CD44+CD62L−CD8+ T cells, and CD44+CD62L+CD8+ T cells in Rabbits immunized with the PPK vaccine (**Fig. 4**). In addition, the PPK immunized rabbits also exhibited higher frequencies of CD4+ CD44+ CD62L+; and CD4+ CD103+ CD69+ cells (**Fig 5**). These results demonstrate that PPK vaccine expressing CXCL11 chemokine and IL-2/IL-15 bolsters the functional HSV-specific CD8+ and CD4+ T cells in the cornea and TG of HLA-A*0201 Tg rabbits.

**Figure 3:**
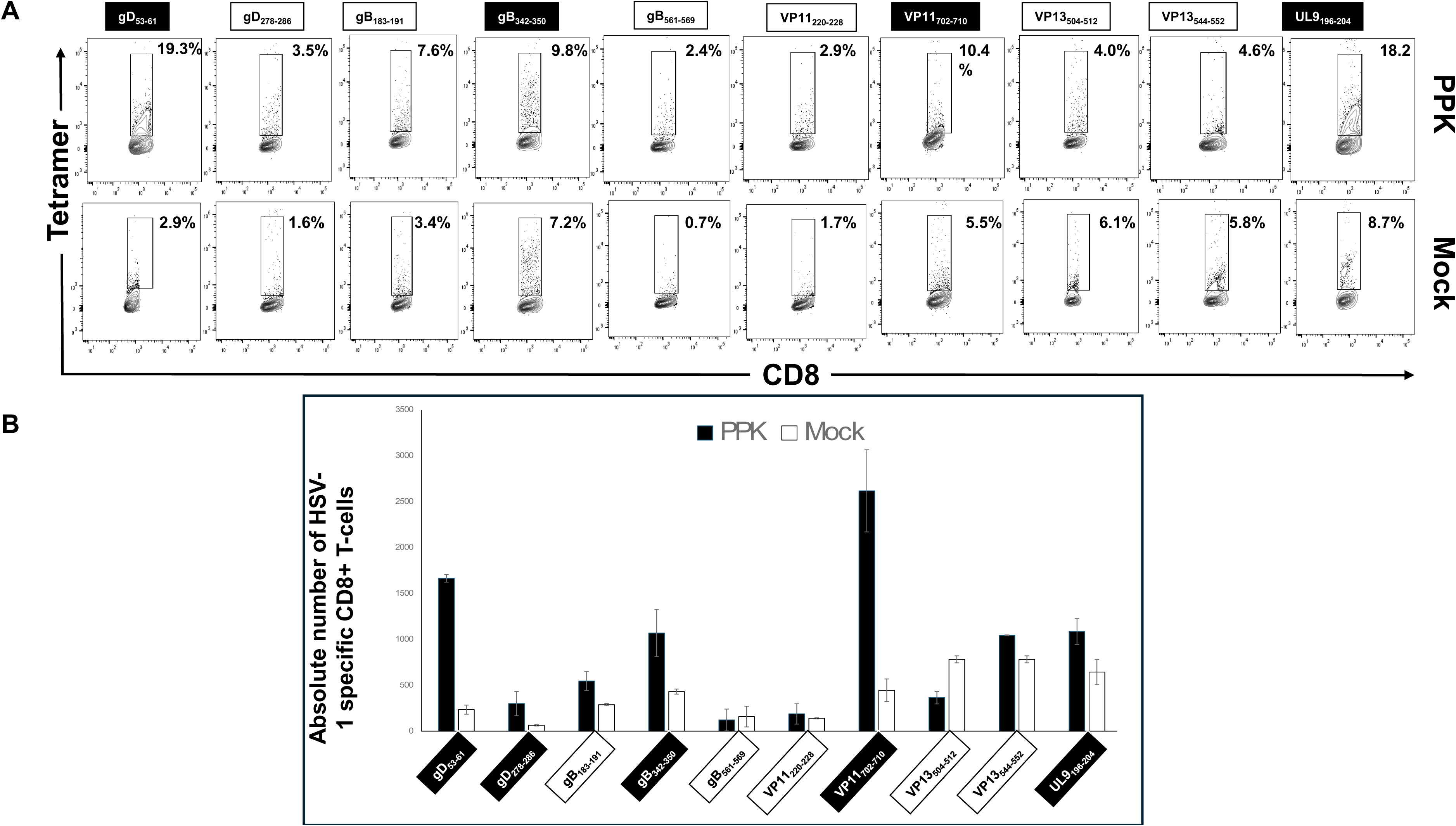
Higher magnitude of HSV-1 specific CD8^+^ T cells observed in PPK vaccinated HLA-A*0201 transgenic rabbits. (**A**) Higher frequencies of CD8^+^ T cells specific to HLA-A*0201-restricted HSV-1 gD_53-61_, gB_342-350_, VP11/12_702-710,_ and UL9_196-204_ epitopes detected by FACS in TG of HLA-Tg rabbits. (**B**) Corresponding bar diagrams showing absolute number of HSV-1 specific CD8+ T-cells among PPK vaccinated and Mock-vaccinated HLA-Tg rabbits. The data are representative of two independent experiments and the graphed values and bars represent SD between the two experiments.

**Figure 4:**
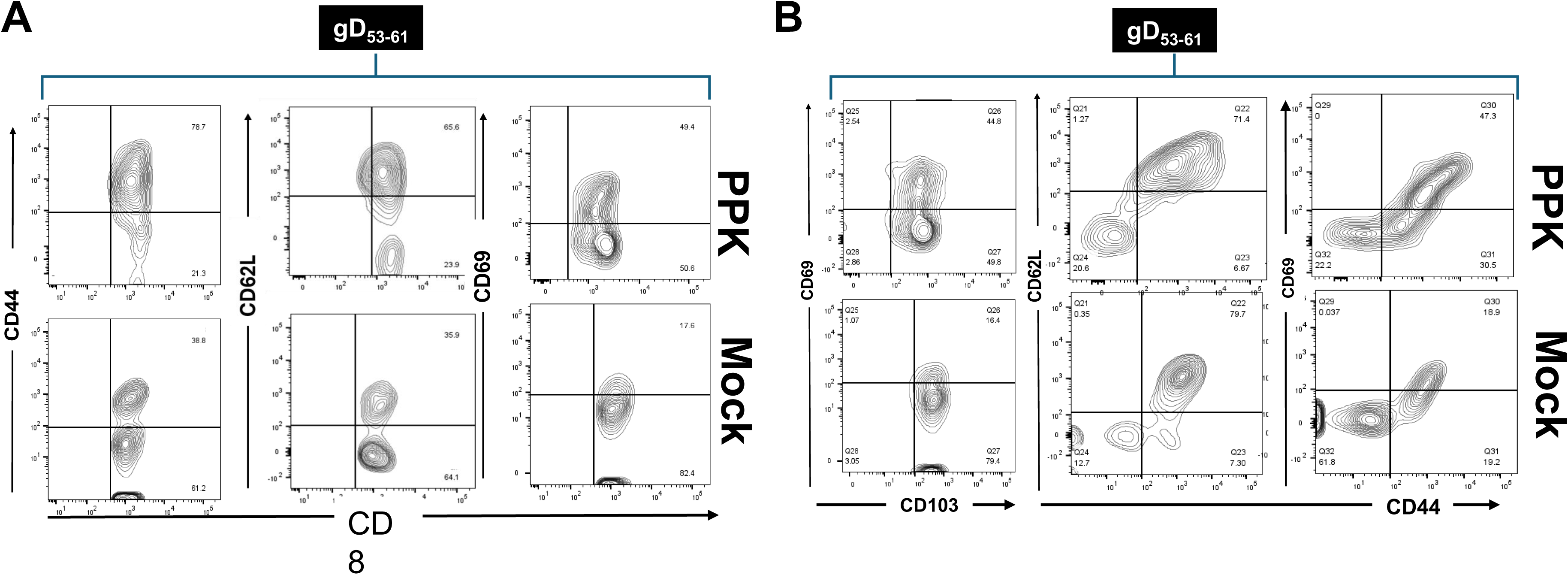
Increased frequencies of CD8^+^ memory T cell population observed in response to Prime-Pull-Keep vaccination approach. (**A-E**) FACS data showing frequencies of memory CD8^+^ T_CM_, CD8^+^ T_EM_, and CD8^+^ T_RM_ cell subsets detected by FACS in HSV-1 infected TG of PPK vaccinated and Mock vaccinated HLA Tg rabbits. To study CD8^+^ T cell memory response in context to PPK vaccination approach multiple markers like CD44, CD69, and CD62L were used.

**Figure 5:**
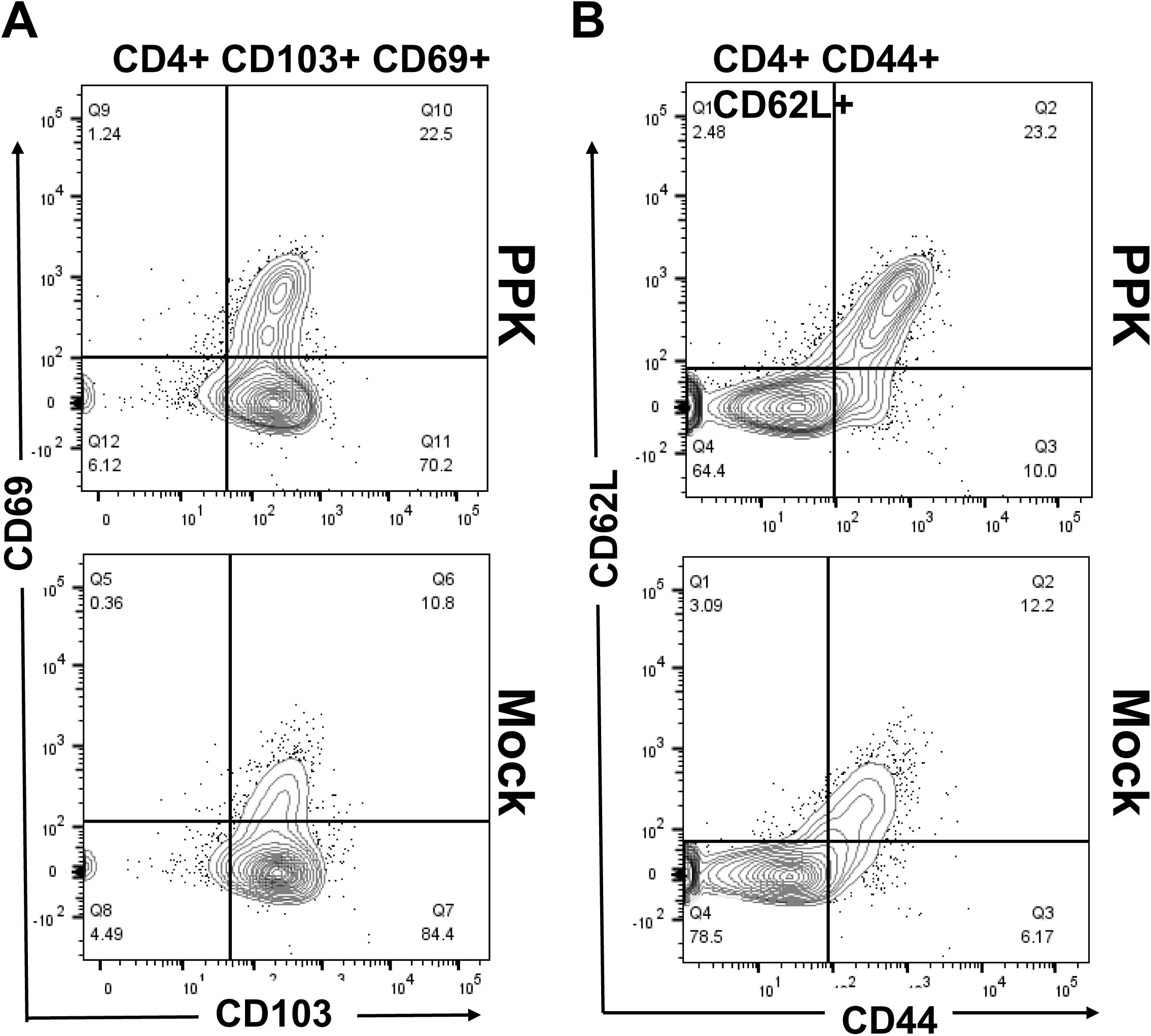
Increased frequencies of CD4^+^ memory T cell population observed in response to Prime-Pull-Keep vaccination approach. (**A-C**) FACS data showing frequencies of memory CD4^+^ T_CM_, CD4^+^ T_EM_, and CD4^+^ T_RM_ cell subsets detected by FACS in HSV-1 infected TG of PPK vaccinated and Mock vaccinated HLA Tg rabbits. To study CD4^+^ T cell memory response in context to PPK vaccination approach multiple markers like CD44, CD69, CD62L, and CD103 were used.

The frequency of HSV-1 specific CD8+ in the spleen as well as in the bloods before vaccination show that both the rabbits groups show no significant difference (**Fig. 6 &7, Supp**). Demonstrating that the PPK vaccine oriented specifically the immune response to the sites of HSV-1 infection TG and cornea. Altogether, these results: (i) indicate that immunization with mixtures of HSV protein specific peptides (Table-1) combined with CXCL11 and IL-2/IL-15 treatment reduced ocular HSV-1 load, and decreased ocular herpetic disease; (ii) suggest that bolstering the number and function of HSV-specific CD8+ T cells that infiltrates the cornea and TG of through a prime/pull/keep vaccine improved protection against ocular herpes infection and disease; (iii) support HLA-A*02:01 Tg rabbits as a useful animal model for investigating the underlying mechanisms by which CD8+ T cells, specific to human HSV-1 CD8+ T cell epitopes, mediate control of ocular herpes infection and disease

**Figure 6:**
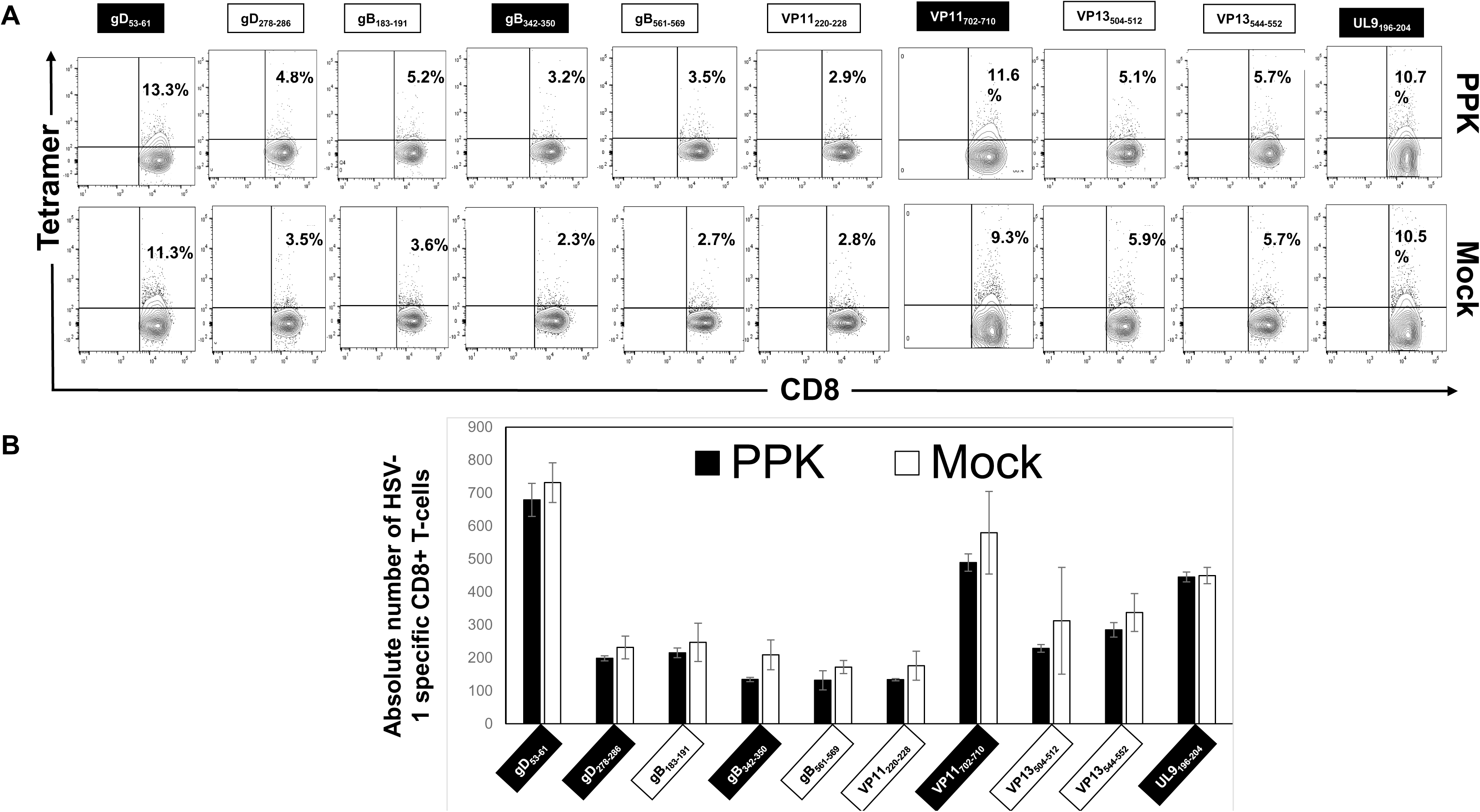
HSV-1 specific CD8^+^ T cells frequencies in PPK vaccinated HLA-A*0201 transgenic rabbits as observed in blood. Data showing frequency of 10 CD8^+^ T cell peptides (gD_53-61_, gD_278-286_, gB_183-191_, gB_342-350_, gB_561-569_, VP11/12_220-228,_ VP11/12_702-710,_ VP13_504-512,_ VP13_544-552,_ and UL9_196-204_) in blood. All HSV-1 infected rabbit showed non-significant differences in HSV-1 specific CD8^+^ T cells frequencies in the blood at day 30 post HSV-1 infection: day 0 before vaccination with PPK or Mock.

**Figure 7:**
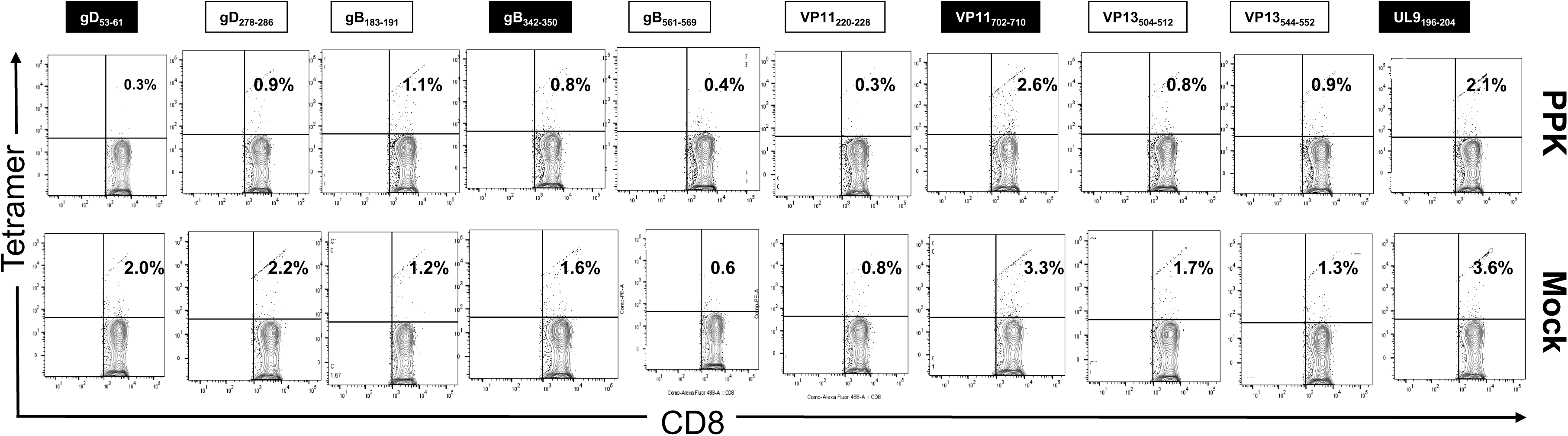
HSV-1 specific CD8^+^ T cells frequencies in PPK vaccinated HLA-A*0201 transgenic rabbits as observed in spleen. Data showing frequency of 10 CD8^+^ T cell peptides (gD_53-61_, gD_278-286_, gB_183-191_, gB_342-350_, gB_561-569_, VP11/12_220-228,_ VP11/12_702-710,_ VP13_504-512,_ VP13_544-552,_ and UL9_196-204_) in blood. All HSV-1 infected rabbit showed non-significant differences in HSV-1 specific CD8^+^ T cells frequencies in the spleen at day 58 post HSV-1 infection, which is also 28 days post vaccination with PPK or Mock

## DISCUSSION

In the present study, we designed multi-epitopes prime/pull/keep HSV-1 Vaccine candidate based on asymptomatic human CD4+ and CD8+ T-cell epitopes as a priming of HSV-1-specific CD4+ and CD8+ cells, CXCL11 for pulling HSV-1-specific CD4+ and CD8+ T cells to the site of infection (TG and Cornea) and IL2 and IL15 to Keep these HSV-1-specific CD4+ and CD8+ T cells in the site of HSV-1 acute and latent infection (TG and Cornea) for longer time (**Fig. 1**). We showed that immunization with a mixture of our PPK vaccine elicited poly-functional CD8^+^ T cell responses in HLA Tg rabbits that was associated with a protection from recurrent ocular herpes infection and disease. Moreover, we observed that immunized HLA Tg rabbits developed frequent and functional HSV-specific CD69^+^CD103^+^CD8^+^ cytotoxic CD8+T cells in the TG and cornea. These potent tissue-resident CD8^+^ T cells protected HLA Tg rabbits from virus reactivation in TG and replication in the eye, and ocular herpetic disease in ocularly challenged rabbits with HSV-1 (McKrae). The results of this pre-clinical trial: (*i*) support implementation of a prime/pull/keep ASYMP peptides-based vaccine strategy to strengthen the protective efficacy of tissue-resident CD8^+^ T cells against ocular herpes; and (*ii*) indicate that ASYMP CD8^+^ T cell epitopes selected from the HSV-1 glycoproteins D and B (gD and gB), viral tegument proteins (VP11/12 and VP13/14) and the DNA replication binding helicase (UL9) proteins are more suitable candidates to be included in the next generation of ocular herpes prime/pull/keep peptide based vaccines.

Complications from the HSV-1 infection range from mild symptoms, such as cold sores and genital lesions, to more serious complications, such as permanent brain damage from encephalitis in adults and neonates and blinding corneal inflammation (3, 5). Ocular HSV-1 infection is the leading cause of viral-induced corneal blindness in industrialized countries. Changes in sexual behavior amongst young adults have been associated with a recent increase in genital HSV-1 infection, resulting from oral-genital rather than genital-genital contact. Approximately 450,000 adults in the United States have a history of recurrent herpetic ocular disease (symptomatic; SYMP individuals), with approximately 20,000 individuals per year experiencing recurrent and potentially blinding ocular herpetic lesions (3, 4, 6, 7). The seropositive SYMP and ASYMP individuals are different with regards to CD8^+^ T cell epitopes-specificity, the magnitude and the phenotype of HSV-specific CD8^+^ T cells (3–7, 18, 29, 31). Thus, a vaccine that converts the presumably non-protective profile of CD8^+^ T cells seen in SYMP patients into the protective profile seen in ASYMP individuals will likely lead to a decrease in ocular herpes. Traditional vaccine formulations using native or recombinant proteins are generally ineffective in the induction of CD8+ T cell responses (32). Clinical trials of HSV vaccines using selected HSV proteins mixed with adjuvant have shown limited efficacy in seronegative women, but not in men (reviewed in (3)). This limitation results from the basic biology of Ag processing and presentation of epitopes to CD8+ T cells, necessitating the endogenous synthesis and presentation of HLA class I molecules. In contrast, our proposed PPK vaccine containing a mixture of the CD4+ and CD8+ T-cell epitopes, CXCL11 and IL2/IL15 induced strong CD8+ T cell responses. In the present study, we demonstrated that immunization with mixtures of peptide vaccines exclusively bearing human epitopes from the HSV 1 glycoproteins D and B (gD and gB), viral tegument proteins (VP11/12 and VP13/14) and the DNA replication binding helicase (UL9) proteins that are mainly recognized by CD8^+^ T cells from HSV-1 seropositive healthy ASYMP individuals and CXCL11 and IL2 and IL15 (1), reduced infectious virus in tears and lessened ocular herpes following ocular challenge in prophylactically immunized HLA Tg rabbits. This peptide vaccine excludes SYMP epitopes that are recognized mostly by CD8^+^ T cells from SYMP individuals with a history of numerous episodes of recurrent ocular herpes disease. We have shown that these ASYMP epitopes in tandem with CXCL-11 and IL1/IL15, PPK vaccine given therapeutically to latently infected HLA Tg rabbits: (*i*) significantly decrease virus reactivation from TG (virus shedding in tears) and/or recurrent ocular disease; and (*ii*) increase the numbers and functions of local HSV 1 glycoproteins D and B (gD and gB), viral tegument proteins (VP11/12 and VP13/14) and the DNA replication binding helicase (UL9) epitopes specific CD8^+^ T cells over the existing immune response induced by the primary infection.

Both rabbit and mouse ocular herpes models have been successful for studying ocular HSV-1 infection and immunity, and each model resulted in new information and discoveries related to human HSV-1 ocular disease (reviewed in (33) and (34)). Mice have been the animal model of choice for most immunologists over the years and results from mice have yielded remarkable insights into the role of CD8^+^ T cells in protection against primary herpes infection (7, 18, 35–38). Unfortunately, spontaneous reactivation of HSV-1 in mice is extremely rare so the relevance of these findings to in vivo HSV-1 spontaneous reactivation cannot be determined in mice (39). The rabbit ocular herpes model has been especially important for investigating viral reactivation and recurrent ocular disease (33, 34, 40). Unlike mouse eyes, but similar to human eyes, the surfaces of the rabbit eyes are relatively immunologically isolated from systemic immune responses (33, 41, 42). Using the “humanized” HLA transgenic rabbit model of ocular HSV-1 that mounts “human-like” CD8^+^ T-cell immune responses (HLA Tg rabbits), we found that immunization of HLA Tg rabbits with the three human CD8^+^ T-cell epitopes induced strong CD8^+^ T cell-dependent protective immunity against ocular herpes infection and disease. From a practical standpoint, the size of rabbit corneas is significantly larger than those of mice and offer a plentiful amount of tissues for phenotypical and functional characterization of HSV-specific T cells using individual tissues (18, 28, 33, 41, 42). To overcome the hurdle of rabbits that do not mount T cell responses specific to human HLA-restricted human epitopes, we introduced the “humanized” HLA Tg rabbit model of ocular herpes whereby the rabbits express human leukocyte antigen (HLA class I molecules) (29, 43). Ocularly-infected HLA Tg rabbit mounted HLA-A*02:01-restricted CD8^+^ T cell responses to HSV-1 glycoproteins D and B (gD and gB), viral tegument proteins (VP11/12 and VP13/14) and the DNA replication binding helicase (UL9) epitopes similar to that from HLA-A*02:01 positive HSV seropositive humans. Since expression of the rabbits’ own MHC class I molecules might interfere with the human HLA-A*02:01-restricted responses (9), and to ensure that all rabbits had a high level of expression of HLA-A*02:01 molecules in over 90% of their CD8**^+^** T-cells, only those HLA Tg rabbits with these characteristics were used in this investigation. These results confirm our previous report that all gD epitopes recognized by CD8^+^ T cells from HSV-1 infected HLA Tg rabbits are recognized by CD8^+^ T cells from HLA-A*02:01 positive HSV seropositive humans (29). Thus, the HLA Tg rabbit is a useful model for pre-clinical testing of candidate vaccines bearing human T-cell epitopes. The HLA Tg rabbit model allows to test the hypothesis that a vaccine that induces appropriate human T-cell responses to HSV-1 can decrease HSV-induced ocular disease. These findings should guide the development of a safe and effective T cell-based herpes vaccine.

In conclusion, four principal findings were determined in the pre-clinical results obtained with HSV-1 infected HLA Tg rabbits in this report. First, a human herpes vaccine that exclusively contains a mixture of human ASYMP CD8^+^ T cell epitopes derived from the HSV-1 gB, gD and VP11/12 proteins along with CXCL11, IL2 and IL15 provides protection in HLA-A*02:01 Tg rabbits against ocular herpes infection and disease. Second, frequent poly-functional HSV-1 ASYMP epitopes-specific CD8^+^ T-cells were induced by the ASYMP human epitopes from gB, gD and VP11/12 proteins and correlated with protection against ocular herpes infection and disease in HLA Tg rabbits, following HSV-1 ocular challenge. Third, the results demonstrate for the first time that the IL2 and IL15 chemokine axis is of paramount importance in keeping the protective CD8^+^ T cells to the TG and corneal tissues associated with clearance of ocular herpes infection and disease for longer period. Fourth, the study validates the HLA-A*02:01 Tg rabbit model of ocular herpes for pre-clinical testing of future herpes vaccine candidates bearing human ASYMP CD8^+^ T cell epitopes against ocular herpes. Overall, the pre-clinical results determined in HLA Tg rabbits draw attention to the prime/pull/keep vaccine strategy, as an alternative to currently used protein-based vaccines, to strengthen the protective efficacy of tissue-resident CD8^+^ T cells against ocular herpes infection and disease.

## ACKNOWLEDGEMENTS

This work is supported by Public Health Service Research R01 Grants EY026103, EY019896 and EY024618 from National Eye Institute (NEI), R21 Grant AI110902 and R41 Grant AI138764-01 from National Institutes of allergy and Infectious Diseases (NIAID) (to L.B.M.), and in part by The Discovery Center for Eye Research (DCER) and the Research to Prevent Blindness (RPB) grant.

This work is dedicated to the memory of late Professor Steven L. Wechsler “Steve” (1948-2016), whose numerous pioneering works on herpes infection and immunity laid the foundation to this line of research. We thank Dale Long from the NIH Tetramer Facility (Emory University, Atlanta, GA) for providing the Tetramers used in this study.

